# Identifying rare, medically-relevant genetic variation in a diverse population: opportunities and pitfalls

**DOI:** 10.1101/2020.05.28.122457

**Authors:** Kevin M. Bowling, Michelle L. Thompson, David E. Gray, James M.J. Lawlor, Kelly Williams, Kelly M. East, Whitley V. Kelley, Irene P. Moss, Devin M. Absher, E. Christopher Partridge, Anna C.E. Hurst, Jeffrey C. Edberg, Gregory S. Barsh, Bruce R. Korf, Gregory M. Cooper

## Abstract

**Purpose:** To evaluate the effectiveness and specificity of population-based genomic screening in Alabama.

**Methods:** The Alabama Genomic Health Initiative (AGHI) has enrolled and evaluated 5,369 participants for the presence of pathogenic/likely pathogenic (P/LP) variants using the Illumina Global Screening Array (GSA), with validation of all P/LP variants via Sanger sequencing in a CLIA-certified laboratory before return of results.

**Results:** Among 131 variants identified by the GSA that were evaluated by Sanger sequencing, 67 (51%) were false positives (FP). For 39 of the 67 FP variants, a benign/likely benign variant was present at or near the targeted P/LP variant. Importantly, African-Americans were significantly enriched for FP variants, likely due to a higher rate of non-targeted alternative alleles close to array-targeted P/LP variants.

**Conclusion:** In AGHI, we have implemented an array-based process to screen for highly penetrant genetic variants in actionable disease genes. We demonstrate the need for clinical validation of array-identified variants in direct-to-consumer or population testing, especially for diverse populations.

## BACKGROUND

Over the past decade, direct to consumer (DTC) genetic testing has become broadly popular amongst people looking to better understand their ancestry or medical health risk. It has been estimated that more than 26 million people in total have ordered some flavor of DTC testing [1, 2]. A recent survey of primary care and specialist physicians found that 35% of respondents report having received a DTC genetic result from a patient [3]. It is also becoming increasingly common for consumers to submit their raw array data to third party websites for interpretation, such as Promethease, codegene.eu, or WeGene [4, 5]. These observations indicate that DTC genetic testing results are increasingly likely to impact clinical decision making by both patients and providers.

Several studies have suggested that DTC genetic testing leads to identification of variants that appear to be medically relevant but fail confirmatory testing and are false positives (FPs)[6, 7]. This is, at least in part, a consequence of the fact that most clinically relevant, disease-associated alleles are rare [8] and, by definition, have low prior probabilities of being present in any given person, particularly within asymptomatic individuals [9]. Further, the majority of DTC testing is conducted using single-nucleotide polymorphism (SNP) arrays, and population-scale studies, such as All of Us [10], are also likely to assay many individuals by SNP arrays. SNP arrays are designed to assign genotypes for commonly observed, rather than rare, variants; they rely upon, for example, empirically observed patterns of genotype-dependent fluorescent intensity clustering rather than absolute assessments of allelic presence or absence [7, 11].

Thus, as the number of tested individuals and interest in clinical genetics grows, it will become important to define the accuracies of SNP-array based detection of highly penetrant variation [6]. FP detection of P/LP alleles is a major concern given the potentially life-altering medical interventions that may follow. Further, false negatives may also be of concern, particularly to the extent that they lead to “false reassurance” in which an individual interprets a negative result to indicate a lack of genetic disease risk [12].

The Alabama Genomic Health Initiative (AGHI) is a state-wide research program to conduct genetic testing for healthy populations, as well as for individuals with rare disorders, to determine utility in disease prevention, management, and treatment. As part of AGHI, we have tested 5,369 (to date) Alabamians in an attempt to identify clinically actionable genetic variation that may be relevant to participant health. This population screen is being conducted using the Illumina Global Screening Array (GSA), which has probes designed to genotype ∼160,000 rare variants. Many of these rare variants were targeted because they reside in clinical disease databases and are annotated as highly penetrant contributors to disease [13, 14]. The AGHI population screen differs from DTC genetic testing as it is free to participants and provides access to genetic counseling, including phone-based or face-to-face counseling for those who receive positive findings. AGHI also provides customized reports of non-informative results to individuals with a personal or family history of disease that is consistent with elevated genetic risk, along with recommendations and resources for follow-up testing. In addition, with participant consent, genetic data are linked to electronic medical records to facilitate future research about the consequences and utility of genetic disease risk assessments.

## METHODS

### Study participant population

Genotyping via AGHI is available to all adult Alabama residents. The research protocol is approved by the UAB Institutional Review Board. Permanent and “pop-up” enrollment sites are located throughout the state to provide access to a broad, diverse population. Participants are recruited via media, social media, and word of mouth. During enrollment, participants meet with a team member to discuss benefits, risks, limitations, and logistics, collect demographic data and contact information, and provide informed consent. Participants can elect to participate in a biobank, have results shared with a healthcare provider, and/or allow re-contact about future research. Two 4 mL blood draws into EDTA tubes are collected; one is used to isolate DNA and the other is archived in a biobank (with consent). Additionally, a health history questionnaire identifies participants with a personal or family history relevant to the medically actionable gene list (described below). This is assessed by genetic counselors to identify participants whose history suggests a genetic risk factor and who should receive recommendation for follow-up testing even if genotyping results are negative.

### Population screen gene list

Variants returned to participants reside within medically actionable genes. These genes were defined by the study team and overlap genes on the ACMG SFv2.0 list [15].

### Genotyping and quality control

DNA isolation was conducted at UAB using the Gentra Puregene blood kit (Qiagen), or at the HudsonAlpha Institute Clinical Services Laboratory (CSL) using the QIAsymphony DSP DNA kit, following standard CAP/CLIA-approved protocols. Genotyping was completed at HudsonAlpha using the Illumina Global Screening Array (GSA-24, v1.0 (1,807 individuals) and GSA-24, v2.0 (3,562 individuals)) per Illumina’s recommended protocol.

Signal intensities detected by the GSA were converted to genotypes using Illumina AutoCall software with a GenCall threshold of 0.15. Primary quality control requirements included per-sample log R standard deviation (SD) less than 0.25 and call rates greater than 98.5% across the array (GSA-24, v1.0), or greater than 99.0% across the autosomes and chromosome X (GSA-24, v2.0).

### Filtration and manual curation

Technical data reports are generated by the CSL and genotype calls are converted to a VCF-like format. Samples are analyzed in batches of various sizes, depending on activity levels at recruiting sites. The number of chromosome X heterozygous counts are calculated per individual to predict sex, and pair-wise kinships values [16] are used to determine participant relatedness. Metrics are tracked to detect potential sample mix-ups, and relatedness is accounted for when defining batch- and study-wide allele frequencies.

GSA-identified variation is filtered to reduce variants to a number that can be manually curated. For each batch, we removed variants that: have a batch allele count >5 (counting alleles within related groups of participants only one time); are present in ExAC/GnomAD [17] or 1000 Genomes [18] at a frequency >0.1%; or are not protein- or splice-site-altering and have a CADD-scaled score below 15 [19]. For *MUTYH* variation, population database frequency thresholds are increased to 2%. Additionally, we filtered variants using a pre-defined list of genes (the ACMG SFv2.0). Although study-wide allele counts, which are accumulating over time, and no-call rates are not used in automated filtration, these values are used during manual curation.

Additionally, batch-level plots of raw fluorescence intensity values are generated for all variants that pass initial filtration, and these plots are used during manual curation.

### Variant classification

Variant classification was conducted using the evidence codes and rules set forth in the American College of Medical Genetics and Genomics (ACMG) guidelines [20].

### Variant validation

Although individual AGHI components are conducted within the HudsonAlpha CSL, a CAP/CLIA-accredited laboratory, the overall workflow is carried out as a research project under an IRB-approved protocol (IRB-170303004). Medically relevant and returnable variants were validated by Sanger and separately interpreted by an independent CAP/CLIA-certified laboratory (PerkinElmer or Emory Genetics Laboratory).

### Return of results

Participants with no genotyping findings receive a non-informative result report via mail. Those who were flagged as being at elevated risk of disease based on personal or family history are given customized reports highlighting their histories and the limitations of AGHI genotyping to detect clinically relevant variants, and are encouraged to seek clinical follow-up with specialists as appropriate. Participants with positive genotyping results receive a phone call from a genetic counselor to describe results, implications, and next steps (face-to-face counseling is available upon request). An individualized research result report based on standard templates is written by genetic counselors and sent to the participant. With consent, a letter is also shared with the participant’s healthcare provider.

### Data Sharing

GSA-identified and Sanger confirmed P/LP variants have been submitted to ClinVar (study ID: AGHI_GT).

### Estimation of continental ancestry

To infer continental ancestry of AGHI participants, we created a continental ancestry reference dataset based on 1000 genomes [21]. To ensure maximum overlap with the GSA, we filtered GSA v2.0 variants using vcftools v0.1.7 to include only autosomal, bi-allelic SNPs and, to reduce LD effects, we thinned SNPs to retain one variant every 2000 bp. We used the isec command of bcftools v1.9 to select the intersecting variants from the 1000 genomes data and vcftools to select variants at a global MAF>5% to create our 1000 genomes ancestry reference dataset. For each batch of GSA samples, we used bcftools to select all variants with at least one heterozygous genotype and that intersect with the ancestry reference dataset. After recoding each set into plink format using plink v1.9.0, we used Admixture v1.3.0 in supervised mode to train an ancestry model on the filtered, superpopulation-labeled 1000 genomes variants (K=5), and then used Admixture v1.3.0 in projection mode to predict the percentages of continental superpopulation ancestry in each of the participants.

## RESULTS

### Population screening in AGHI via GSA

To date, we have enrolled, genotyped, and analyzed 5,369 individuals, 75% of which are female. 73% self-report as European American, 20% as African-American, and 2% as Asian. The remaining 5% represent individuals of Native American/Alaskan Native or Native Hawaiian/Pacific Islander descent, or their race is unknown. 3.3% are Hispanic/Latino, and 45% live in medically underserved areas [22].

We have used the Illumina GSA to genotype AGHI participants. While the GSA mostly targets sites of common variation, a subset of probes (∼160,000 of 654,027) are designed to target rare, potentially clinically relevant genetic variation. Many of these variants were included on the array because they reside in clinical genetics databases like ClinVar [13, 14], and thus may be highly penetrant contributors to “actionable” diseases. Currently in AGHI, we return only P/LP variants in clinically actionable disease genes predominantly associated with cancer and cardiac risk (see Methods).

Variants of interest are filtered and selected based on a number of attributes, manually curated, classified according to ACMG criteria, and Sanger validated prior to return to study participants (Figure 1). Personal and family history of disease is asked of each individual at time of enrollment. While variants are classified in accordance with ACMG guidelines [20], family history information is available to the study team.

**Figure 1:**
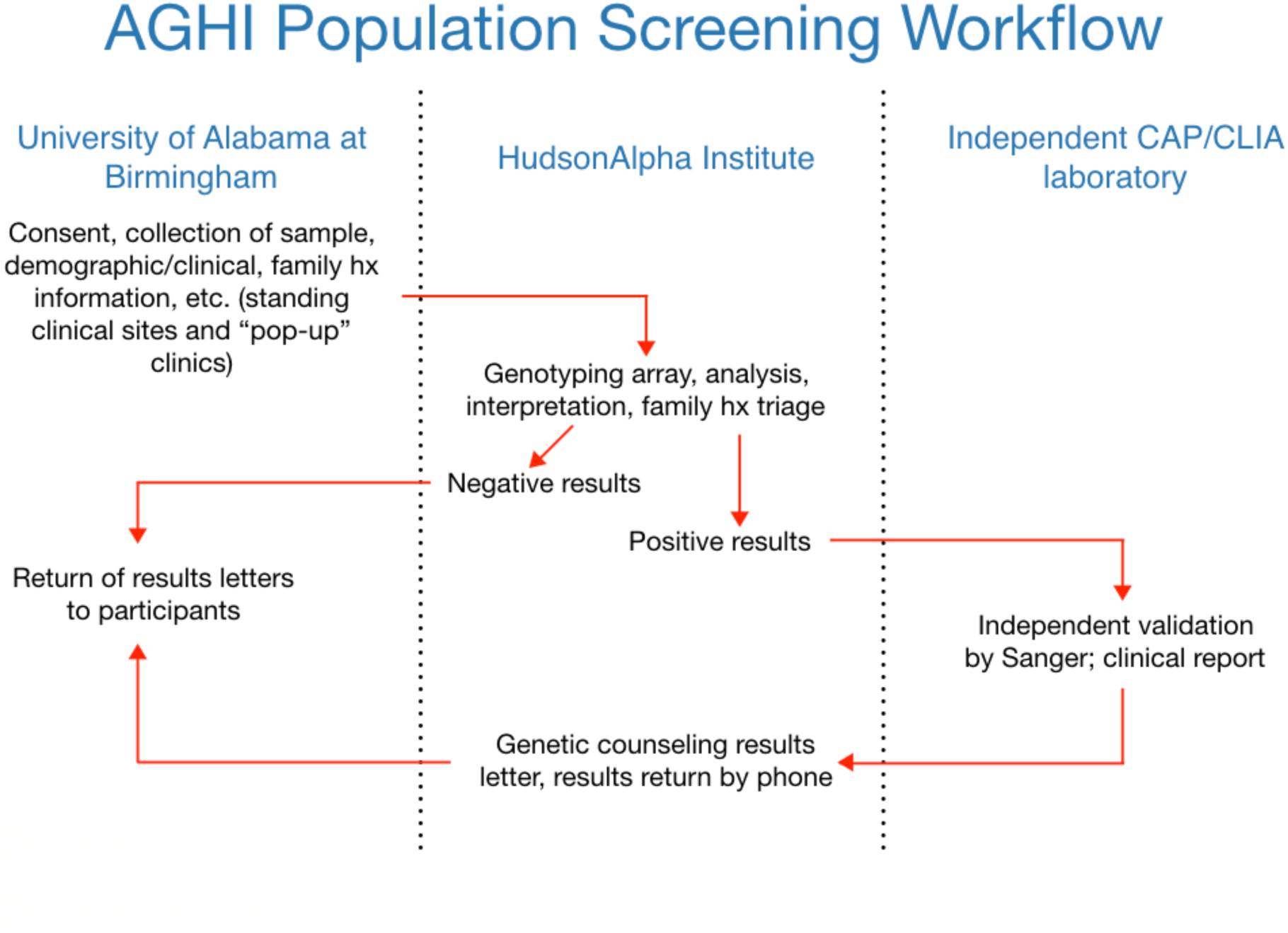
Workflow of population screening in the Alabama Genomic Health Initiative.

### Sensitivity of the GSA to medically relevant variants

To assess sensitivity of the GSA prior to conducting population screening in AGHI, we compared variation targeted by the GSA to “secondary finding” variants found via genome or exome sequencing of 7,422 healthy individuals as reported in previous studies [23, 24]. Among 126 unique P/LP secondary findings, 57% are targeted by v1.0 of the array (71% by v2.0), providing an estimate of the upper limit on sensitivity (Table S1). Most (62%) P/LP secondary finding variants found by sequencing that are not targeted by the GSA (v2.0) are rare loss-of-function (LOF) alleles (nonsense, frameshift, or canonical splice site). This reflects the fact that LOF variants that are not in clinical genetic databases can often meet ACMG criteria for P/LP status [20]. Missense alleles not previously seen in affected individuals typically do not meet P/LP criteria due to ambiguity associated with missense variation.

To experimentally assess sensitivity of the array to targeted P/LP variants, we tested DNA samples from 20 previously sequenced individuals (85% European ancestry) known to harbor at least one variant targeted by the GSA (23 unique variants, 15 genes; all heterozygous). All expected variants were identified correctly by the GSA, suggesting high sensitivity among array-targeted variants (Table S2).

### True positive findings in AGHI

To date, 5,369 individuals from the population cohort have undergone GSA testing and analysis. To confirm array-detected variation, we conducted Sanger testing for 131 unique variants found to be heterozygous (with exception of one homozygote) among 191 individuals, comprising 204 total person-by-variant Sanger tests (Table S3). Of the 131 variants tested, 99 were tested in only one individual, and among these 48 were confirmed while 51 were not. Among the 30 variants tested in multiple individuals, 14 confirmed in all tested individuals and 16 failed in all tested individuals, while one variant confirmed in 1 of 2 individuals and one variant confirmed in 1 of 5 tested individuals. Overall, 62 variants validated in all individuals tested (47%), 67 did not validate in any individual tested (51%), and two validated in a subset of tested samples.

Among the variants that validated in at least one individual, 57 lie within a medically actionable gene, were classified as P/LP, and were returned to study participants. Overall, 80 of 5,369 total individuals were confirmed to harbor at least one clinically actionable P/LP variant, resulting in an overall yield of 1.5% (one individual harbors two findings). Returned variants reside in 19 different genes, including nine unique variants in *BRCA2* (11 individuals), seven in *BRCA1* (nine individuals), and five in *MYBPC3* (nine individuals; Table 1). Of the 62 variants that validated in all tested individuals, 10 are predicted to result in frameshift (16%), 15 result in nonsense (24%), 29 result in missense substitution (47%), seven are predicted to alter splicing (11%), and one leads to in-frame deletion (2%, Figure 2).

**Table 1:** Returnable P/LP genetic variation was detected by GSA, and Sanger validated, in 80 AGHI participants across 19 different genes (one individual harbors two findings).

| Gene | Disease association (MIM#) | Unique variants | Number participants (% participants) |
| --- | --- | --- | --- |
| APOB | FHCL2 (144010) | 1 | 6 (0.11) |
| BRCA1 | BROVCA1 (604370) | 7 | 9 (0.17) |
| BRCA2 | BROVCA2 (612555) | 9 | 11 (0.21) |
| GLA | Fabry disease (310500) | 2 | 2 (0.04) |
| KCNH2 | LQT2 (613688) | 1 | 2 (0.04) |
| KCNQ1 | LQT1 (192500) | 3 | 3 (0.06) |
| LDLR | FHCL1 (143890) | 2 | 3 (0.06) |
| MLH1 | HNPCC2 (609310) | 3 | 3 (0.06) |
| MSH2 | HNPCC1 (120435) | 1 | 1 (0.02) |
| MSH6 | HNPCC5 (614350) | 2 | 3 (0.06) |
| MUTYH | FAP2 (608456) | 5 | 5 (0.09) |
| MYBPC3 | CMH4 (115197) | 5 | 9 (0.17) |
| MYH7 | CMH1 (192600) | 3 | 5 (0.09) |
| PKP2 | ARVD9 (609040) | 3 | 3 (0.06) |
| PMS2 | HNPCC4 (614337) | 2 | 2 (0.04) |
| RET | FMTC (155240) | 2 | 2 (0.04) |
| RYR1 | MHS1 (145600) | 3 | 9 (0.17) |
| SCN5A | LQT3 (603830) | 2 | 2 (0.04) |
| SDHB | PGL4 (115310) | 1 | 1 (0.02) |

**Figure 2:**
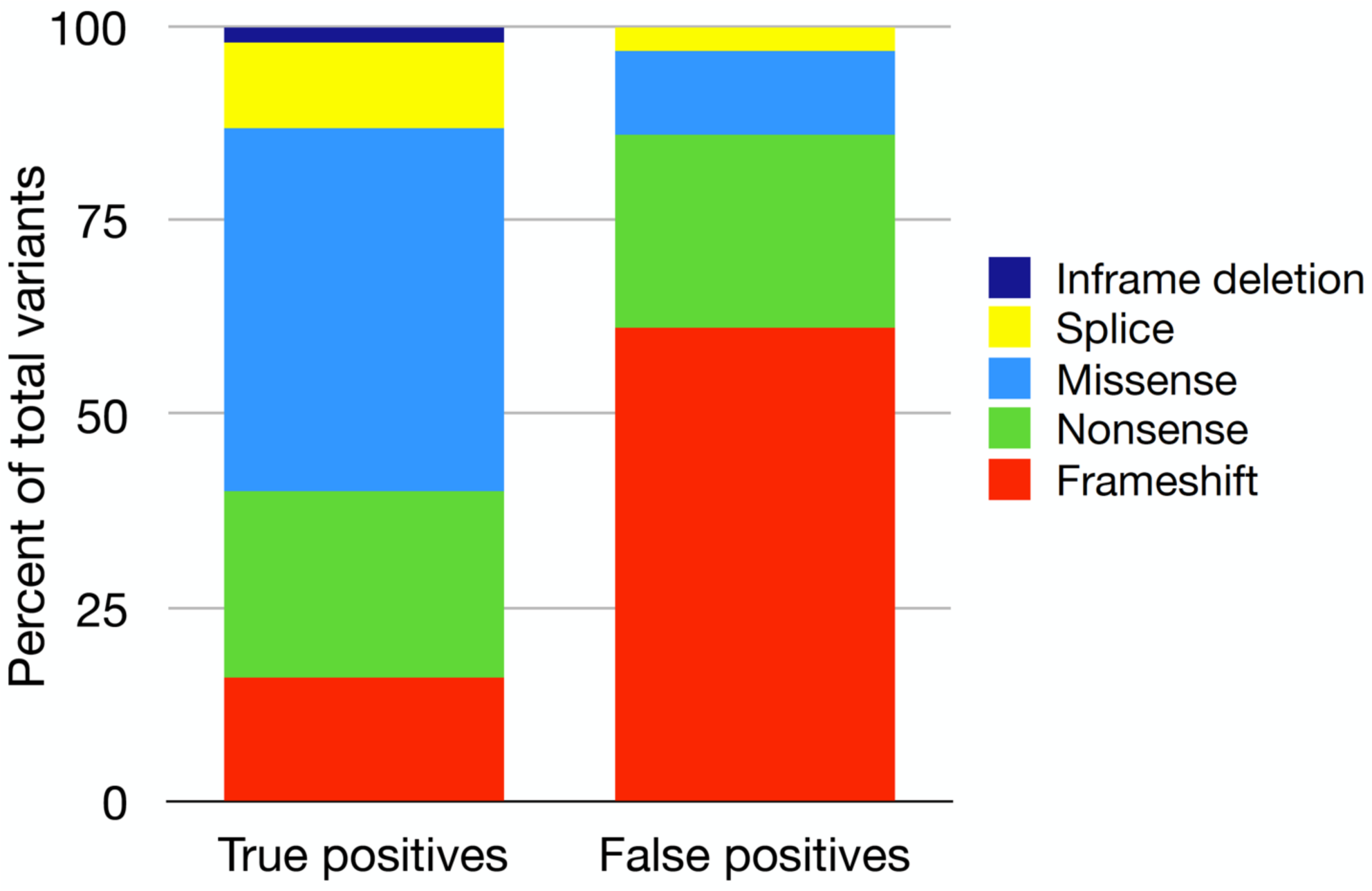
Differences in genetic variation types between true and false positive variants. GSA-targeted variants determined to be false positive by Sanger testing are enriched for failed detection of targeted indels. 61% of false positive variants are predicted to result in frameshift, compared to 16% of true positive events.

### False positive findings

Analysis of GSA data for the first batches of AGHI participants revealed an unreasonably high rate of variants that passed initial QC and were suspected to be P/LP variants in study-defined medically actionable genes. For example, analysis of the first 55 individuals resulted in 13 unique variants across 15 participants (2 pairs of first-degree relatives harbored the same variant) that passed genotype quality and variant filters and were candidates for return. If all these variants were accurately detected, that would suggest a medically relevant variant yield of 27%, a value sharply higher than that seen in other large studies of non-ascertained populations, which tend to be 1-3% even when using sequencing [23-25]. Tellingly, eight of these 13 array-detected variants were classified as P/LP and were Sanger tested, but none were confirmed.

Here, we describe one variant in detail as an illustrative example of the types of errors that result from the use of the GSA in this context. Specifically, NM_000059.3(BRCA2):c.4258delG (p.Asp1420fs), a pathogenic frameshift, was detected in two first-degree relatives within the first 55 participants. Sanger testing of both individuals revealed that they actually harbor NM_000059.3(BRCA2):c.4258G>T (p.Asp1420Tyr), a benign missense variant. Thus, the array detected heterozygosity for a non-targeted benign allele at the targeted genomic position. Across all participants in AGHI, the frameshift variant NM_000059.3(BRCA2):c.4258delG has been reported by the GSA as heterozygous at a frequency of 0.64% among called individuals, an observation that is implausibly high relative to its allele frequency of 0.0008% in gnomAD [17]. In contrast, the benign missense variant (p.Asp1420Tyr) that was Sanger-detected has a frequency of 0.66% in gnomAD. This non-targeted allele is thus likely leading to all of the FP frameshift alleles being flagged by the GSA.

More generally, among the 67 GSA-detected variants that did not validate in any individual, Sanger sequencing detected a benign/likely benign alternative allele at, or near, the tested position in 58% of cases (39/67; Table 2). Further, as part of our sensitivity testing (see above), we also tested two previously sequenced individuals known to have heterozygous variation at a position targeted by the GSA but harboring a non-targeted alternate allele. For both of these variants, the GSA reported that the sample was heterozygous for the array-targeted, rather than the actually present, allele (Table S2), further confirming the effects of non-targeted alleles at targeted positions. These results demonstrate that GSA-detected rare heterozygotes often result from the existence of a non-targeted allele. As the non-targeted alleles are often more common than the targeted alleles and more likely to be benign, this substantially inflates FP detection of disease-associated variation.

**Table 2:**
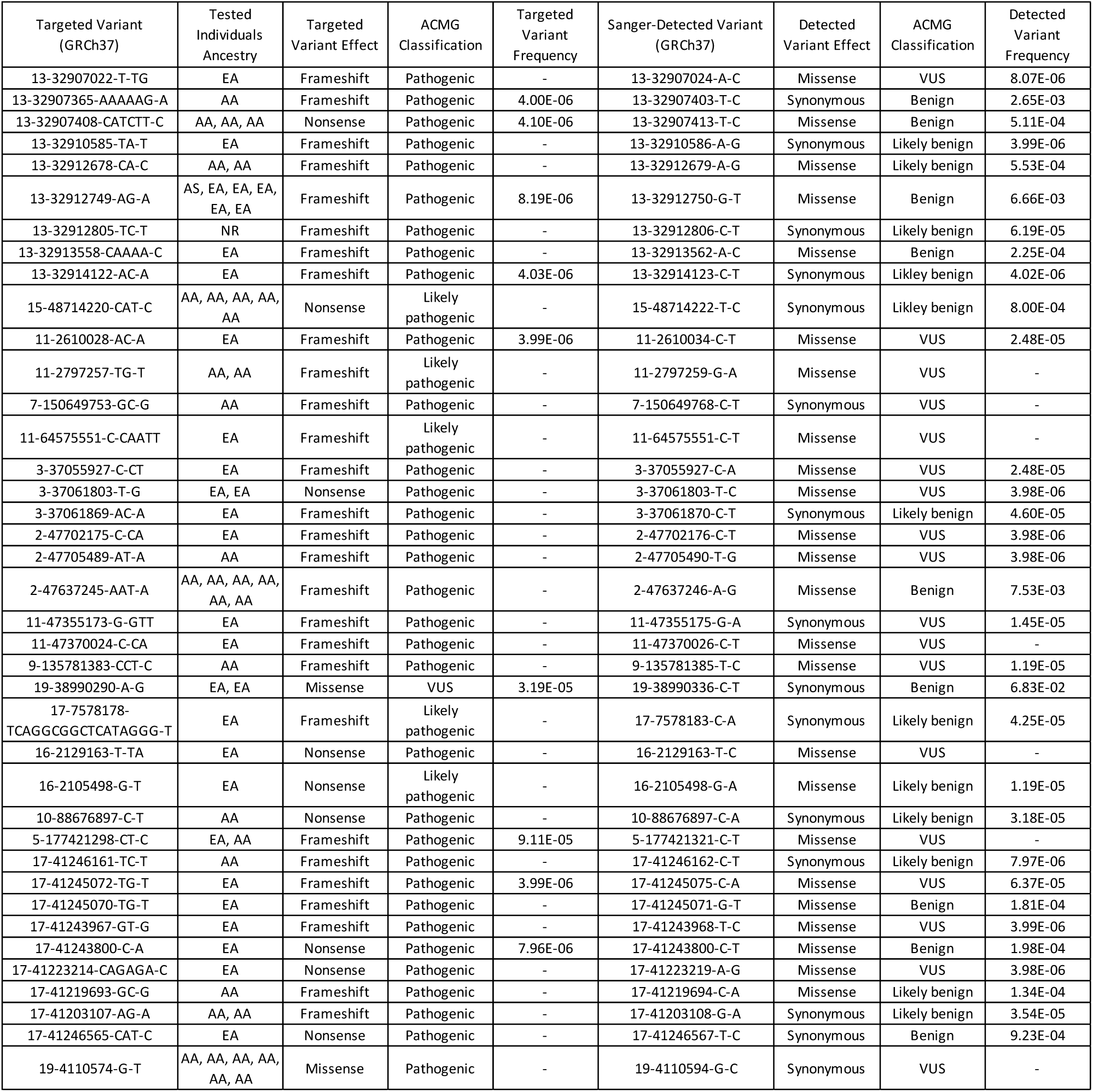
Clinically relevant genetic variation detected by GSA in AGHI participants that did not validate via Sanger testing, but for which Sanger did detect an alternative non-targeted variant. The Sanger-detected variants often represent benign/likely benign missense or synonymous events, in contrast to the P/LP variants targeted by the array (EA = European American, AA = African American, AS = Asian, NR = not reported).

Among the 67 array-targeted FPs, 41 are predicted to result in frameshift (61%), 17 result in nonsense (25%), seven in missense substitution (11%), and two are predicted to alter splicing (3%, Figure 2). In contrast to the predicted effects of the 62 TPs that validated in all individuals tested, FPs are enriched for variation predicted to result in frameshift (61% in FPs versus 16% in TPs), suggesting that most FPs result from failed detection of targeted indels. In contrast, array-targeted missense variation resulting from single-nucleotide variation are more likely to be TPs, at least for the variants that we Sanger tested (11% in FPs versus 47% in TPs).

We examined the other attributes of array-targeted FPs to see whether automated predictors of errors could be determined. We found that variant-level no-call rates are higher (p=0.003) in FPs (0.3%) than in TPs (0.076%), as are study-wide allele frequencies (0.1% in FPs vs 0.02% in TPs; Table S3). Also, we hypothesized that probe uniqueness/mappability may play some part in FP detection. However, when we aligned probes on the GSA to the reference assembly (GRCh37), we found no difference in the number of matching sequences in the genome when comparing FPs and TPs (using both 80% and 90% sequence identity cutoffs; Table S3).

Thus, no-call rates, batch- and study-level allele frequencies, and the presence of known alternative nearby variation are the features that we have found to be most strongly predictive of TP/FP status with respect to medically relevant variation. It should be noted that internal allele frequency and no-call rates have become more effective with accumulation of data as the study progresses, and we increasingly add to the list of variants (such as the *BRCA2* variant mentioned above) that are unreliable and do not warrant curation or Sanger confirmation (although see below). However, we continue to observe a substantial rate of FPs, with 46% of tested, unique variants failing Sanger confirmation among the most recent 1,000 participants.

### False positive findings in the context of race/ethnicity

We have collected self-reported race/ethnicity for almost 98% of enrolled study participants, and have also calculated from GSA data the percentage of continental ancestry for each individual using 1000 genomes phase 3 [21] (see Methods). We assessed error rates in relation to genetic ancestry. We focus here on European ancestry (EA) and African American (AA) individuals, as the total numbers of participants (361) and Sanger tested individuals (10) that are neither EA nor AA is too small to facilitate robust conclusions. Of the combined total 99 participants who self-reported as either EA or AA individuals and underwent Sanger testing which did not validate, 46 of these individuals are AA (46%). This is in contrast to the 22% of all EA/AA participants being AA. Similarly, 22% of the 79 EA/AA individuals harboring GSA-detected genetic variation that did validate self-report as AA. The enrichment of FPs among AAs is substantial (OR=3.2) and statistically significant (p=8e-4; Figure 3). Further supporting this ancestry-correlated FP rate, for the two variants that each validated in one individual but not in others (one failed individual for one variant, and four for the other), the individuals in which the variants validated are EA, whereas those harboring FP variation are AA. Self-reported ancestries correlated strongly with genetically inferred degrees of African continental ancestry, with Sanger-tested individuals who self-reported as AA having an estimated 40-90% African ancestry and the Sanger-tested self-reported EA individuals all being <11% African ancestry (median 1.9%, Table S4).

**Figure 3:**
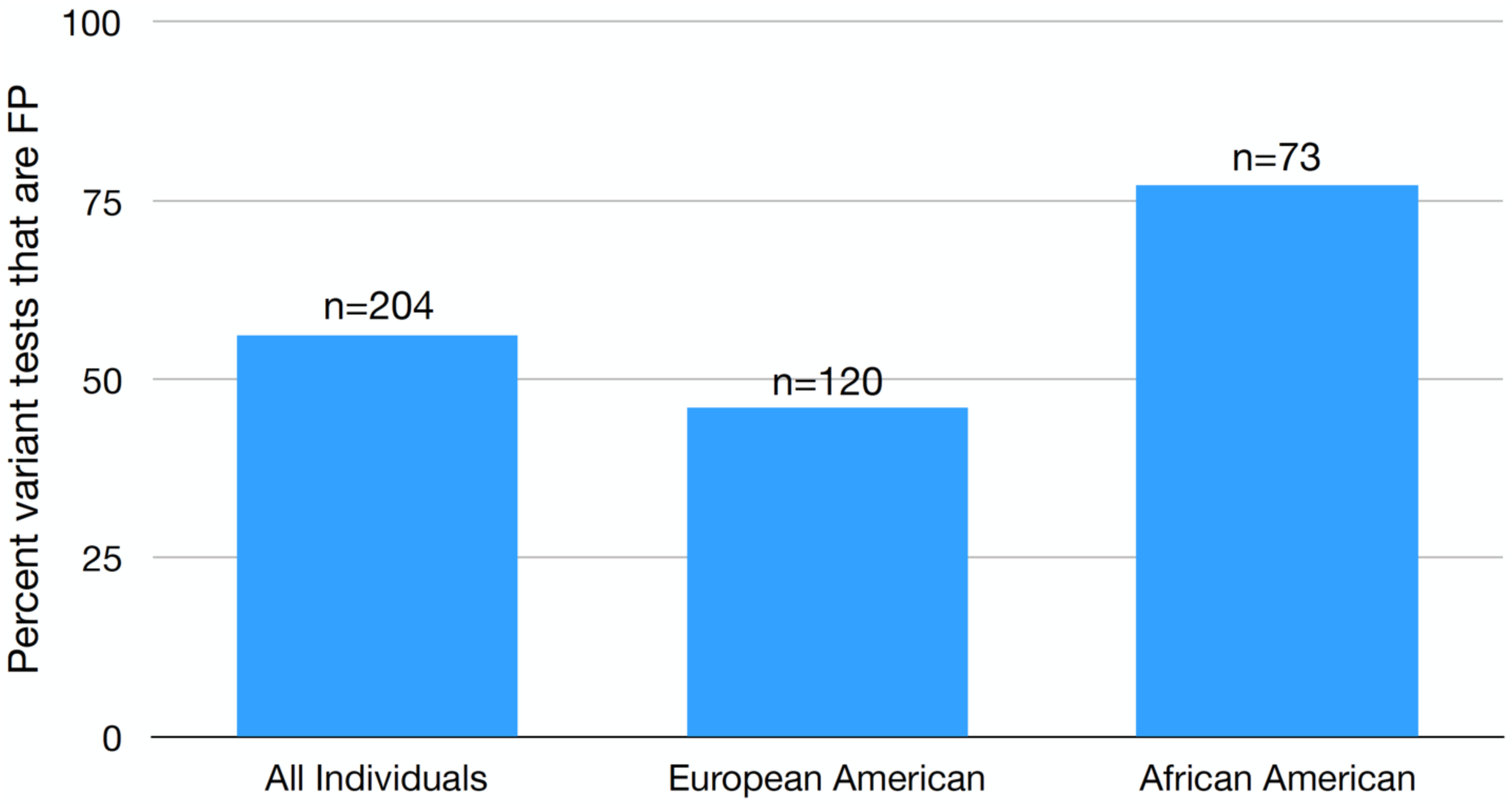
False positive findings in the context of race/ethnicity. False positive genetic variation is more often identified in individuals of African descent. 77% of GSA-detected variants in African American individuals represent false positive calls, in contrast to 46% in individuals of European descent, and 56% across all individuals tested.

## DISCUSSION

To date, 5,369 participants have been genotyped and analyzed in the AGHI population screen to detect P/LP variants in clinically actionable genes. 80 individuals (∼1.5%) were found to harbor a P/LP variant in a subset of medically actionable disease genes associated with cancer or cardiac risk. All P/LP variants identified via the genotyping array and returned to participants were validated by Sanger sequencing in a CAP/CLIA-certified lab to ensure technical accuracy prior to return.

SNP arrays are primarily designed to assign genotypes at sites of common variation. They rely upon empirical genotype-dependent clustering of fluorescence intensity values from many samples [11]. Identification of rare variants by array is thus an exercise in outlier detection (e.g., an individual sample whose fluorescence intensity at a given variant deviates from all other samples in a given batch or, in many cases, from all other samples ever tested in a given study) and is prone to inferring P/LP variants that are false discoveries. Because all variants of interest are Sanger sequenced for confirmation, we can measure array specificity among the population of variants that are classified as P/LP and pass manual quality curation. We have found that many (51%) rare and potentially P/LP genetic variants identified by array do not validate by Sanger, and represent FPs. While we demonstrate that some FPs can be systematically filtered as a result of no-call rates, elevated study and batch-specific allele frequencies, and overlap with known alternative variants, a high rate of FP detection remains even after aggressive filtration and manual curation. For example, 46% of tests have failed validation over the past 1,000 AGHI participants despite removal of variants found to be problematic among the first 4,369 samples.

We have shown that most FPs (58%) result from detection of an allele different from that which is targeted. In many such cases, the array is targeting a rare indel that results in frameshift, while the actual detected alleles often lead to a benign missense/synonymous substitution. Given that more than 61% of non-confirmed variants represent targeted indels, this type of variation is particularly problematic when using array technologies.

We have also provided data relevant to assessing the sensitivity of the GSA to highly penetrant rare variation. We found that most (57% GSA v1.0, 71% GSA v2.0) P/LP “secondary findings” reported in two large sequencing-based studies are targeted by the GSA, and we showed that 23 known variants in 20 samples were all successfully flagged as heterozygotes by the GSA. While these estimates suggest high sensitivity, and support the utility of array-based testing in the context, they overstate actual sensitivity. In particular, the presence of non-targeted, benign alleles near targeted, pathogenic alleles lead to FPs at a subset of targeted P/LP variants. While such P/LP variants are on the array and potentially could be accurately detected in individuals which harbor them, filtration of these calls is needed to curtail an unsustainable rate of FPs, necessarily reducing sensitivity. In fact, we have now defined a list of 67 GSA-targeted P/LP rare variants that cannot be relied upon (Table S3); while small relative to the scope of all rare GSA variants (∼160,000), this is a large fraction relative to all variants flagged for interpretation or follow-up (e.g., 67 unreliable vs 64 total returned unique variants in AGHI).

We also found that ancestry associates with the likelihood that a rare variant is correctly genotyped, with African-Americans being substantially more likely to harbor FPs. Other studies have shown that there are ancestry-associated discrepancies in accurate clinical interpretation of sequencing data, a result that, at least in part, reflects reduced representation of non-European individuals in clinical and research genetic databases [26, 27]. Our results show that ancestry discrepancies also affect the technical quality of array-based detection of rare variants. We note that this is in addition to potential discrepancies in accurate clinical interpretation. Further, while the existence of non-targeted benign alleles near targeted P/LP alleles is an issue across all ethnicities, it is likely true that the reduced representation of non-European alleles makes probe design for the detection of P/LP alleles in non-Europeans less effective; that is, to the extent that probe design accounts for the existence of known alternative alleles [28], such designs will necessarily be less effective when there are fewer known variants within a given population.

Overall, results from screening a large population of Alabamians indicates that FP detection rates among array-based rare variant genotypes are considerable, but manageable. This work supports the findings of others suggesting that array-detected rare genetic variants often represent FPs. Tandy et. al., 2018 [6] report that 40% of clinically relevant variants identified via DTC are FPs, while Weedon et. al. demonstrates that only 16% of array-identified rare variants confirm in sequencing data [7]. Though our results support the hypothesis that population-scale genotyping can detect many individuals at elevated risk for actionable diseases, they also provide clearer definitions of testing limitations. Further, our observations strongly support the conclusion that all array-detected P/LP variants should be confirmed in a clinical genetics laboratory prior to return to patients and clinical providers, especially for individuals of under-represented minority groups.

## ACKNOWLEDGEMENTS

Thank you to all AGHI participants for their contributions to this study. This study was conducted at the University of Alabama at Birmingham and the HudsonAlpha Institute for Biotechnology and funded by the state of Alabama.

## SUPPLEMENTAL TABLES

Table S1: 126 P/LP variants previously identified by genome or exome sequencing used to determine array sensitivity. These variants were identified across 7,422 healthy individuals. 57% of the variants are targeted by GSA v1.0, and 71% by GSA v2.0.

Table S2: Genetic variation previously identified by genome or exome sequencing that are targeted and detected by GSA.

Table S3: All GSA-detected genetic variation identified in the Alabama Genomic Health Initiative screen that was sent for Sanger testing (AC = allele count, AN = allele number, AF = allele frequency).

Table S4: Estimates of African ancestry for AGHI individuals who received Sanger testing.

